# Comparative Genomics of Transisthmian Damselfishes (*Abudefduf saxatilis* and *A. troschelii*)

**DOI:** 10.1101/2025.07.17.665425

**Authors:** Claire B. Tracy, Carlos F. Arias, Eirlys Tysall, Marc P. Hoeppner, W. Owen McMillan, Oscar Puebla, Moisés A. Bernal

**Author notes:** Author for Correspondence: Claire B. Tracy, Department of Biological Sciences, Auburn University, Auburn, AL, USA.

## Abstract

The uplift of the Central American Isthmus (CAI) represents a natural laboratory for the study of allopatric speciation in marine organisms. Several geminate species pairs of both vertebrates and invertebrates formed following the uplift of the CAI, including damselfishes of the Pomacentridae family. However, to date no studies have explored the genomic differences among geminate species in the Pacific and Atlantic Oceans. In this study we present genome assemblies for the transisthmian species pair consisting of the Sergeant Major *Abudefduf saxatilis* (Tropical Atlantic) and the Panamic Sergeant Major *Abudefduf troschelii* (Tropical Eastern Pacific) derived from PacBio long-read sequencing. The new genomes are near-chromosome level, and among the highest-quality genomes currently available for coral reef fishes. We show that large structural variants distinguish the two species, including a 9 megabase inversion in linkage group (putative chromosome) 6. Additionally, we show through an analysis of demographic history that alleles within the two genomes have different coalescence time distributions, which may be due to different effective population sizes, population structures, and/or selection regimes in the two oceans. Finally, we highlight gene families that were significantly expanded or contracted between *A. troschelii* and *A. saxatilis*. Some of these are related to the environmental conditions that differ between the two oceans, such as gamma crystallin M (*crygm*), which is linked to vision, and vitellogenin (*vtg*), which is associated with egg provisioning. These genomes set the stage for comparative analyses of genetic structure and selection on the marine organisms that originated with the formation of the CAI.

**Significance Statement:** Understanding speciation in oceans has represented a historical challenge, given the high rates of dispersal during larval stages for most marine groups. There are few geologic events that have propelled marine biodiversity while completely blocking the genetic exchange during the speciation process. The final closure of the Central American Isthmus (CAI), 2.8 million years ago, is one of these few examples where marine populations from the tropical Pacific and Atlantic oceans have been completely isolated, providing the opportunity to study how genomes diverge in the absence of gene flow. Here, we present the first comparative genomics study of marine organisms separated by the CAI, showing that genomes on the two sides of the CAI differ structurally. We also identify contractions and expansions of various gene families between the two species, which are potentially associated with adaptations to the different environmental conditions in the two oceans. This represents a relevant first step for comparative genomics of geminate species isolated by a major geologic event.

## Introduction

The uplift of the Central American Isthmus (CAI) is a major geological event that led to speciation in many marine taxa. This occurred over millions of years as landmasses gradually uplifted via tectonic movements, leading initially to the formation of islands (the Panama Arc) throughout the Central American Seaway, and making oceanic connections between present day Pacific and Atlantic oceans shallower (O’Dea et al., 2016). This process caused gradual changes in the biotic and abiotic environments that built up over time and culminated in the final closure of the CAI approximately 2.8 million years ago (MYA) (O’Dea et al., 2016) causing the complete isolation of the Tropical Pacific Ocean from the Tropical Atlantic Ocean.

This physical closure caused the cessation of gene flow between the two ocean basins, while simultaneously causing major shifts in ocean currents, resulting in starkly different environments in the Tropical Eastern Pacific Ocean (TEP) and Tropical Western Atlantic Ocean (TWA). The TEP experienced an increase in productivity due to the uplift of cooler waters driven by trade winds, compared to the TWA which experienced a reduction in upwellings changing nutrient availability and causing the Caribbean to become warmer, more saline, and less productive (Jackson & O’Dea, 2023). These changes have been shown to also have biotic effects, decreasing presence of suspension feeders and predators in the Caribbean (Jackson & O’Dea, 2023). Today several areas of the TEP are still characterized by seasonal upwellings (i.e. January through April; D’Croz & O’Dea, 2007; O’Dea et al., 2012). In contrast, most of the TWA is characterized by warmer waters and oligotrophic conditions year-round, with only a few small areas of upwelling as the exceptions (D’Croz & O’Dea, 2007; Jackson & O’Dea, 2023; Rueda-Roa & Muller-Karger, 2013). The formation of the CAI is recognized as one of the most consequential events in marine biogeography, representing a natural laboratory for studying the processes of genetic divergence, speciation, and adaptation in marine species.

This vicariant event led to the formation of many geminate species pairs (sister species separated by the formation of a barrier; Jordan, 1908). While the timing of the uplift and final closure of the CAI has been debated, with some estimates of final closure closer to 10-30 MYA (Bacon et al., 2015; Montes et al., 2015), an estimate of final closure at 2.8 MYA is now largely accepted (O’Dea et al., 2016). This closure, and the formation of geminate species pairs, has been the topic of extensive research. A large body of literature has sought to elucidate the exact timing of species divergence to better understand whether speciation occurred upon final closure, or whether it occurred earlier throughout the millions of years of gradual uplift leading up to the final closure (Lessios 2008). As a result, it is now generally understood that the timing of divergence can vary substantially among taxa. Some species have diverged prior to the final closure of the CAI, including molluscs (*Arca mutabilis & A. imbricata*; > 30MYA; (Marko, 2002)), fishes (Eleotridae, Apogonidae, 5-15 MYA; (Thacker, 2017) and Haemulidae; (Tavera et al., 2012)), and snapping shrimps (genus *Alpheus*; 3-18 MYA; (Hurt et al., 2009; Knowlton & Weigt, 1998)). These instances were likely a result of restriction of gene flow due to the gradual uplift of islands and a presumed change in depth across the Central American Seaway. Fewer molecular studies have evaluated the potential for reproductive isolation among species on either side of the CAI. These few examples include shrimps (Knowlton et al., 1993) and sea urchins (Lessios, 1984; Lessios & Cunningham, 1990), and in many cases hybridization is still possible despite millions of years of separation (Lessios, 2008). Despite these advances, questions remain on the role geographic isolation and adaptation to local conditions have played in the differentiation of species separated by the CAI at the genome level.

One pair of geminate taxa that was formed by the uplift of the CAI are Sergeant Majors (damselfishes, Pomacentridae), with the Panamic Sergeant Major (*Abudefduf troschelii*) in the TEP and the Sergeant Major (*Abudefduf saxatilis*) in the Tropical Atlantic (Figure 1). We note that these two species are not geminate species strictly speaking since *A. saxatilis* is closely related to the African Sergeant *A. hoefleri* in the Eastern Atlantic, and this divergence process appears to have taken place after the divergence from *A. troschelii* (Campbell et al., 2018; Tang et al., 2021). To reflect this, we do not refer to *A. saxatilis* and *A. troschelii* as geminate species but transisthmian species instead. Estimates date the divergence between *A. troschelii* and *A. saxatilis* around 2.4 MYA to 2.5 MYA, coinciding with the final closure of the CAI at 2.8 MYA (Campbell et al., 2018; McCord et al., 2021; Quenouille et al., 2004; Rabosky et al., 2018; Tang et al., 2021). Today, *A. saxatilis* is distributed throughout the tropical and subtropical Atlantic Ocean, in warm, tropical waters (Fishelson, 1970), and *A. troschelii* is more narrowly distributed throughout the TEP and experiences colder and more variable temperatures throughout its range (Robertson, 2009; Figure 1). Prior to their divergence, the common ancestor was likely distributed throughout the Central American Seaway, and the uplift of the CAI formed two isolated populations that underwent allopatric speciation. The different environments experienced by this transisthmian species pair provide an opportunity to evaluate the role of geographic isolation and adaptation in the divergence of closely related species.

**Fig. 1.**
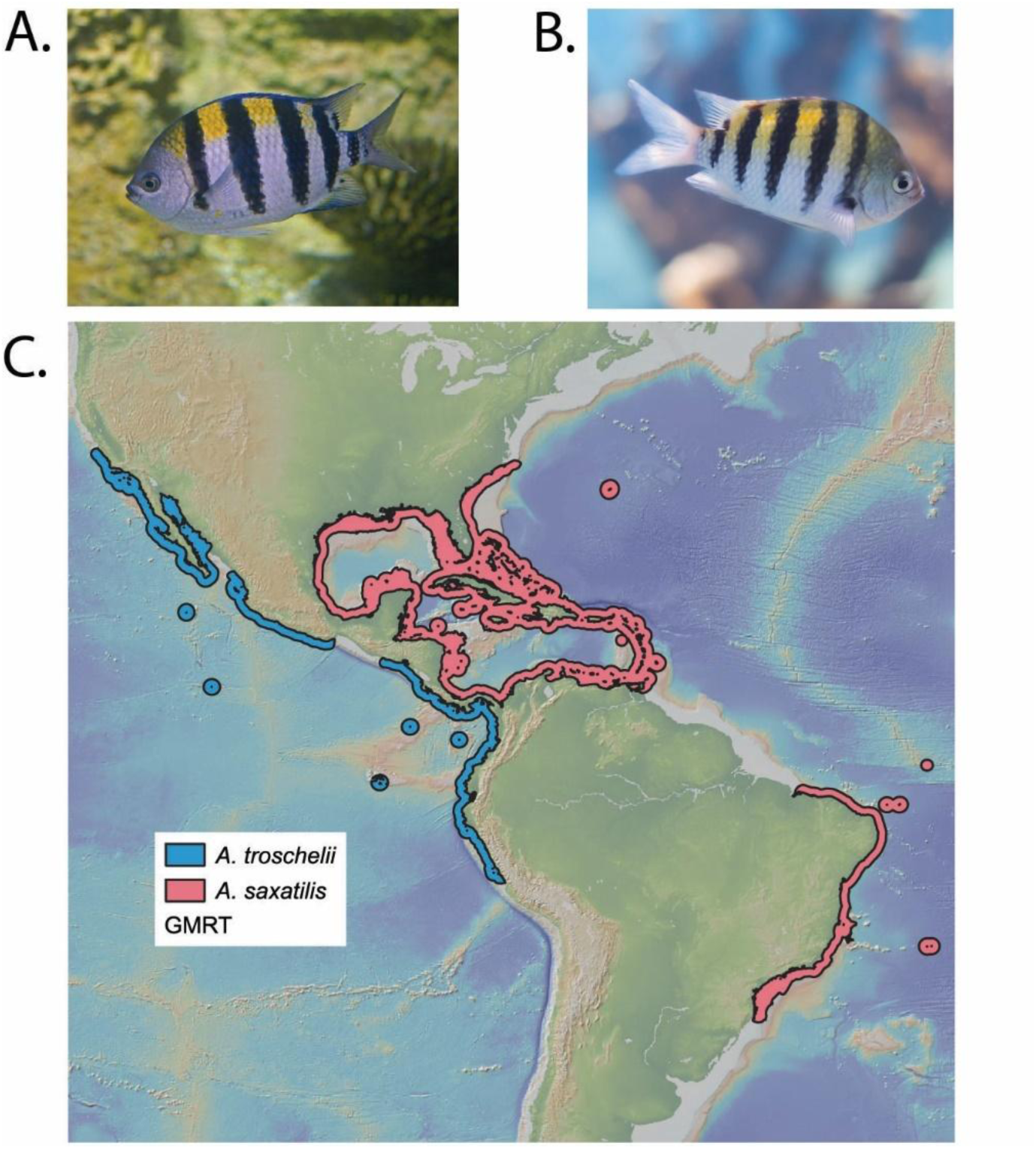
Images and new-world distributions of the two species considered in this study. (A) Image of *Abudefduf troschelii*, image credit: Hectonichus, License CC-BY-SA 3.0, (B) Image of *Abudefduf saxatilis*, image credit: Matthew T Rader, MatthewTRader.com, License CC-BY-SA 4.0. (C) Map of the distribution of *A. troschelii* (blue) and *A. saxatilis* (pink) limited to their new-world distributions, obtained from the IUCN RedList.

Despite decades of molecular analyses of geminate species separated by the CAI (Knowlton & Weigt, 1998; Lessios, 2008; Thacker, 2017), few studies have leveraged whole-genome sequencing to study their evolutionary history. Long-read technologies increase the reliability of de novo assemblies in non-model taxa, leading to higher quality reference genomes for subsequent analysis of population divergence, gene expression, and epigenetics. Long-read sequences are particularly well suited for the assembly of long repetitive regions, and to identify large structural variants (SVs) by spanning these regions—including SV breakpoints—entirely (Luan et al., 2020; Mérot et al., 2020). This is relevant as structural variants can play a role in speciation and have been shown to be associated with environmental adaptation in both plants and animals (Akopyan et al., 2022; Cayuela et al., 2021; Fuller et al., 2018; Kirkpatrick, 2010; Mahmoud et al., 2019; Tigano et al., 2018; Zhang et al., 2021).

This study presents the first long-read genomes for a transisthmian species pair, and the first genomes for the genus *Abudefduf*. These fishes were chosen based on their role in coastal food webs, abundance in shallow marine ecosystems, as well as their territorial behavior, representing a group of great ecological relevance in tropical coastal regions of the Americas (James Cooper et al., 2009; Villegas-Hernández et al., 2022). Within this group the *A. troschelii*/*A. saxatilis* pair was selected because it most likely diverged close to the final closure of the CAI. The main aims of this study were to: 1-compare the structure of the genomes of a transisthmian species pair; 2-compare the historical effective population sizes of the two species; and 3-identify gene families that have expanded or contracted during the differentiation of the two species. This study represents a first step forward in understanding the complex impact of the rise of the CAI on the divergence of marine organisms, while also developing new genomic resources for the study of damselfishes.

## Results and Discussion

### Genome Assembly and Annotation

The genome of one individual of *A. saxatilis* from Florida was sequenced using PacBio. The resulting de novo genome assembly had a total length of 811 megabases (Mb) across 287 contigs with an N50 of 25.8 Mb, a maximum contig length of 36 Mb, a minimum contig length of 13 kilobases (Kb), a GC content of 40.96%, a heterozygous read coverage of 44X, and a homozygous read coverage of 88X (Table 1). The heterozygosity was determined to be 0.563% (Figure S1) which is similar to previously reported measures of heterozygosity in other teleost species (Tigano et al., 2021). The Benchmarking Universal Single-Copy Orthologs (BUSCO) analysis of genome completeness identified 3,598 (98.85%) complete genes, 30 missing genes (0.82%) and 12 (0.33%) partial genes.

**Table 1.**
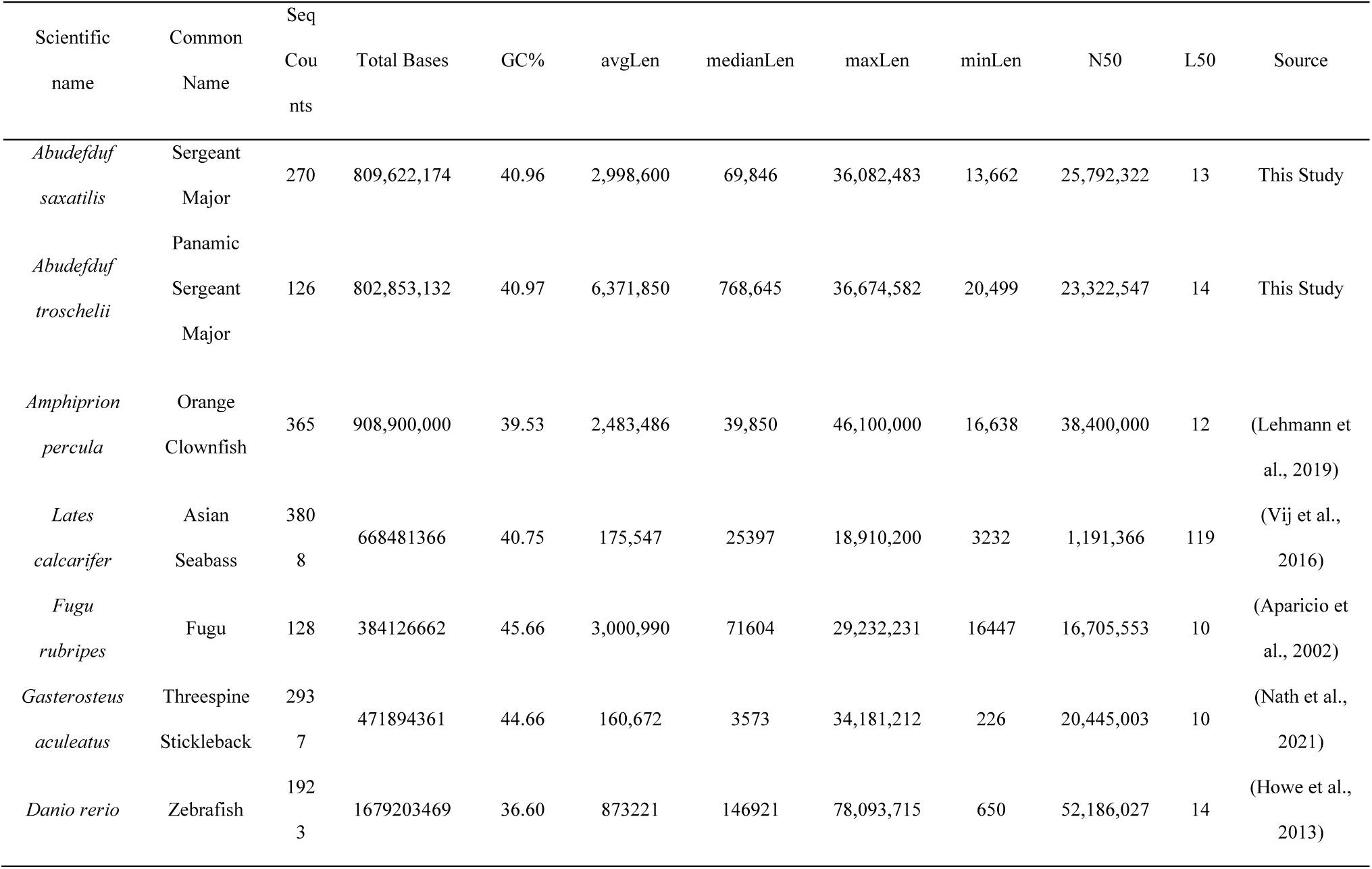
Assembly statistics of the de novo assembly of *A. saxatilis* and *A. troschelii* compared to previously published and annotated fish genomes.

The genome of one individual of *A. troschelii* from Panama was also sequenced using PacBio. The de novo genome assembly of *A. troschelii* had a total length of 802 Mb across 126 contigs with an N50 of 23 Mb, a heterozygous read coverage of 20X, and a homozygous read coverage of 39X (Table 1). The heterozygosity was determined to be 0.593% (Figure S1). The BUSCO analysis identified 3,595 (98.76%) complete genes, 32 missing genes (0.88%) and 13 (0.36%) partial genes. Both genomes had higher completeness than many other published teleost genomes to date (Table 2). We additionally examined how raw reads mapped back to the assembled genomes using Merqury (Rhie et al., 2020). We found a consensus quality value (QV) of Q63.88 and a k-mer completeness of 90.68% for *A. troschelii*, and a QV of Q61.08 and a k-Mer completeness of 91.83% for *A. saxatilis* (Figures S2-S3).

**Table 2.**
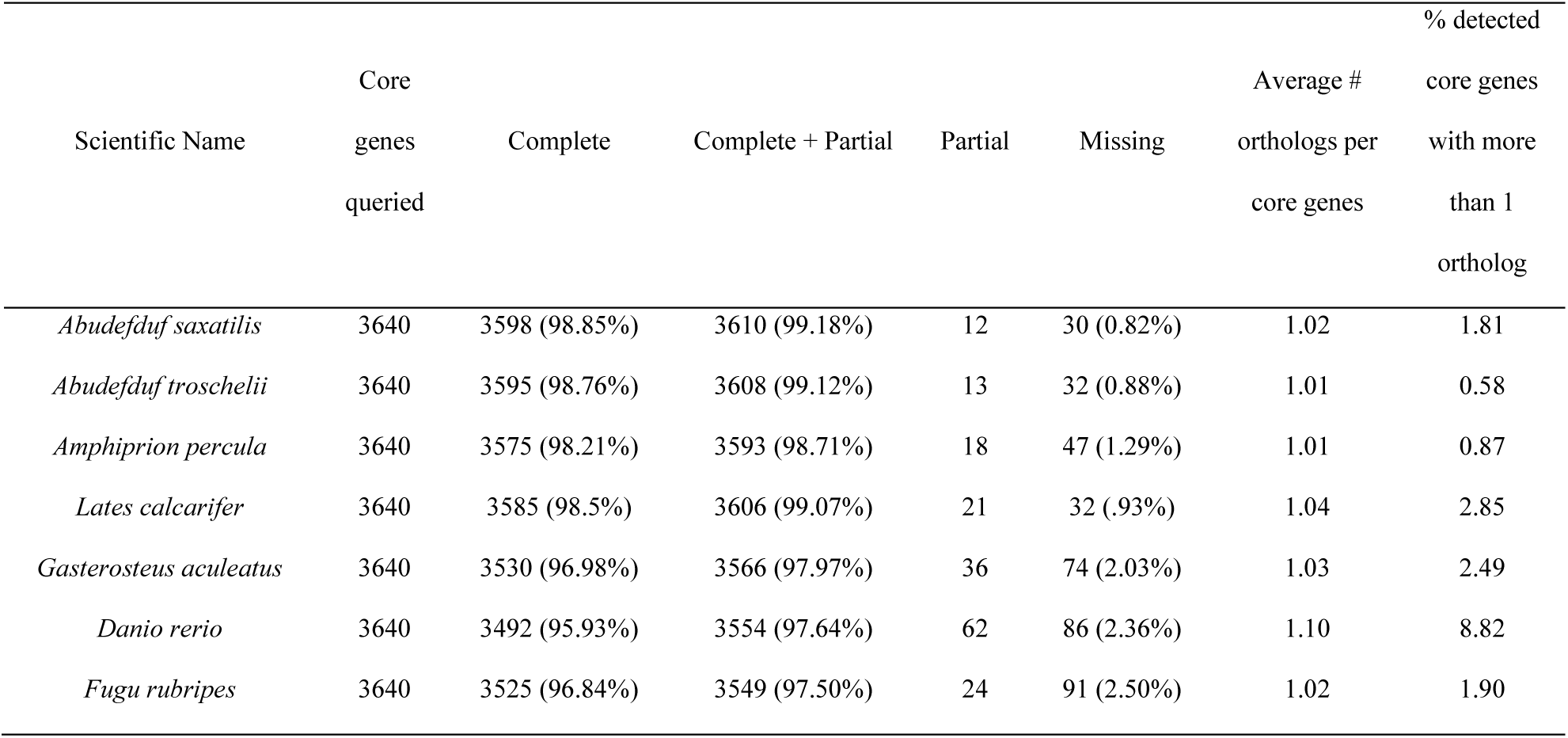
BUSCO statistics detailing the completeness of the genome of *A. saxatilis* and *A. troschelii* compared to previously published and annotated fish genomes. All statistics based on BUSCO v. 5 to enable comparison among genomes.

### Synteny

The de novo assemblies of both species produced large contigs that were near chromosome length. To confirm the contig lengths and compare them to the known chromosomes of another damselfish with a chromosome-level genome available, the genome of *A. saxatilis* was mapped to the genome of the orange clownfish (*Amphiprion percula*). Sixteen chromosomes from the *Amphiprion percula* assembly were covered by 1-4 contigs from the *A. saxatilis* assembly, while 8 chromosomes were covered by 8-31 contigs from the *A. saxatilis* assembly (Table S1).

As expected, there is high synteny between *A. saxatilis* and *A. troschelii*, with large blocks of synteny and some inversions or chromosomal rearrangements between the two (Figure 2). Three large potential inversions were identified (Figure 2): a 28 Mb inversion hereafter called Inversion 1 (*A. saxatilis*: ptg000023; *A. troschelii*: ptg000042); an 8 Mb inversion called Inversion 2 (*A. saxatilis*: ptg000026; *A. troschelii*: ptg000001); and another 8 Mb inversion termed Inversion 3 (*A. saxatilis*: ptg000019; *A. troschelii*: ptg000019).

**Fig. 2.**
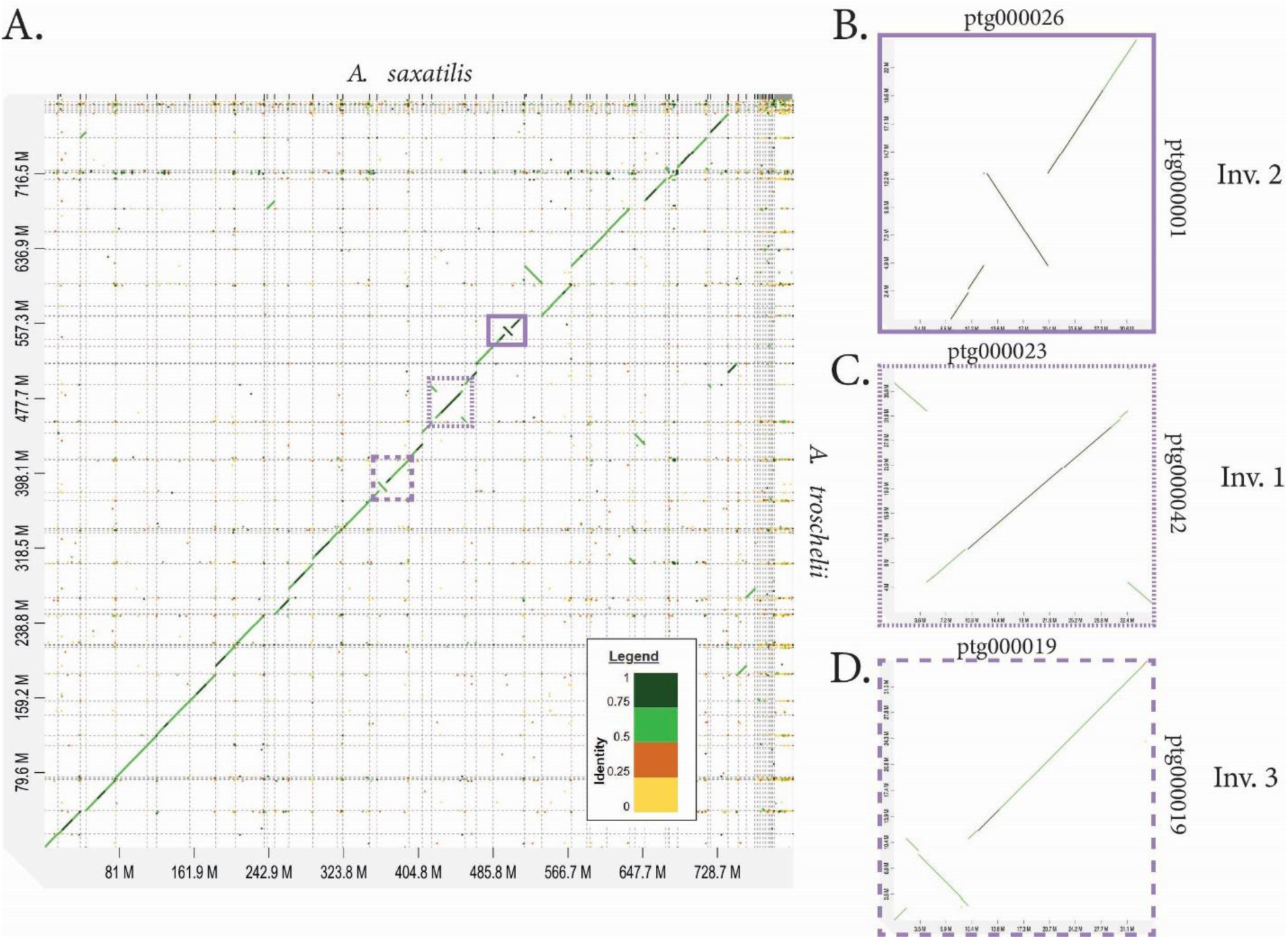
Analysis of synteny mapping the *A. saxatilis* assembly (x-axis) to the *A. troschelii* assembly (y-axis) to visualize large rearrangements or structural variants between the two genomes. Genes falling along the diagonal are matches to one another, and potential structural rearrangements are seen when genes fall off the diagonal or when the diagonal is in the opposite direction. (A) Synteny analysis between the entire genome of *A. saxatilis* (x-axis) and *A. troschelii* (y-axis) from D-GENIES. The analysis shows three potential inversions across the genome shown in panels B-D.

We manually examined the genes around the inversion breakpoints (5 genes before, first and last 5 within inversion, and 5 after) for both species. Around the Inversion 1 breakpoints, genes that stood out include gonadotropin releasing hormone receptor (*GNRHR*) and three high choriolytic enzyme 1-like genes in *A. saxatilis*, however these genes were not annotated in our *A. troschelii* genome. No notable genes seeming to relate to the environment stood out around the Inversion 2 breakpoints. Meanwhile, around the Inversion 3 breakpoints, genes that stood out include empty spiracles 3 homeobox (*emx3*), and fibroblast growth factor receptor-like 1 (*fgfrl1*) in *A. saxatilis*, while fork-head box (*FOXl1*) was found in *A. troschelii*.

A search for specific gene ontology (GO) categories that may be over-represented in each inversion compared to the rest of the genome found no significant genes above the 10% false discovery rate (FDR) for any of the three possible GO categories (Molecular Function, Biological Process, or Cellular Component) for *A. saxatilis* in any of the three inversions, or for *A. troschelii* on Inversion 1. For Inversion 2 in *A. troschelii* one significant GO category above the 10% FDR was detected for MF (4/8 amine transmembrane transporter activity, p-value = 0.006), BP (4/8 amine transmembrane transporter activity, p-value = 0.006), and CC (7/65 brush border membrane, p-value = 0.095). The amine transmembrane transporters allow amines to be transported across a membrane which could include neurotransmitters such as serotonin, dopamine, and norepinephrine impacting synapses (Purves et al., 2001). The analysis on *A. troschelii* Inversion 3 found one significant GO category above the 10% FDR for BP (activation of transmembrane receptor protein tyrosine kinase activity, p=0.009) and MF (transmembrane receptor protein tyrosine kinase activator activity, p=0.032), but none were found in CC. These transmembrane receptor tyrosine kinases play a role in cell signaling and growth (E. Li & Hristova, 2010).

Inversions have been associated with the potential for local adaptation (Akopyan et al., 2022; Kirkpatrick, 2010). However, to make a better assessment of the roles of inversions in the divergence of the two species, whole genome sequencing within and across populations of each species will be required to determine the prevalence and age of the inversions, and whether they have implications for local adaptation. Determining the age of the inversions will also be crucial for understanding the timing of speciation, and whether speciation may have occurred allopatrically prior to the final closure of the CAI.

### Demographic History

The results of the Pairwise Sequentially Markovian Coalescence (PSMC; H. Li & Durbin, 2011) analysis are presented in Figure 3. The first thing to note is that the analysis fails to go back until the divergence point between the two species, potentially due to a scarcity of alleles with deep coalescence times in *A. troschelii*. This illustrates the limitation of this approach when it comes to addressing the initial divergence of transisthmian species, even when it occurred upon final closure of the CAI. Nevertheless, our objective was not to date the divergence time between the two species with the PSMC analysis since such an estimate would rely heavily on mutation rate and generation time estimates, which are uncertain. In this regard divergence estimates provided by phylogenetic methods, in particular the ones that are independent of the timing of the final closure of the CAI, appear more reliable (Campbell et al., 2018; McCord et al., 2021; Quenouille et al., 2004; Rabosky et al., 2018; Tang et al., 2021).

**Fig. 3.**
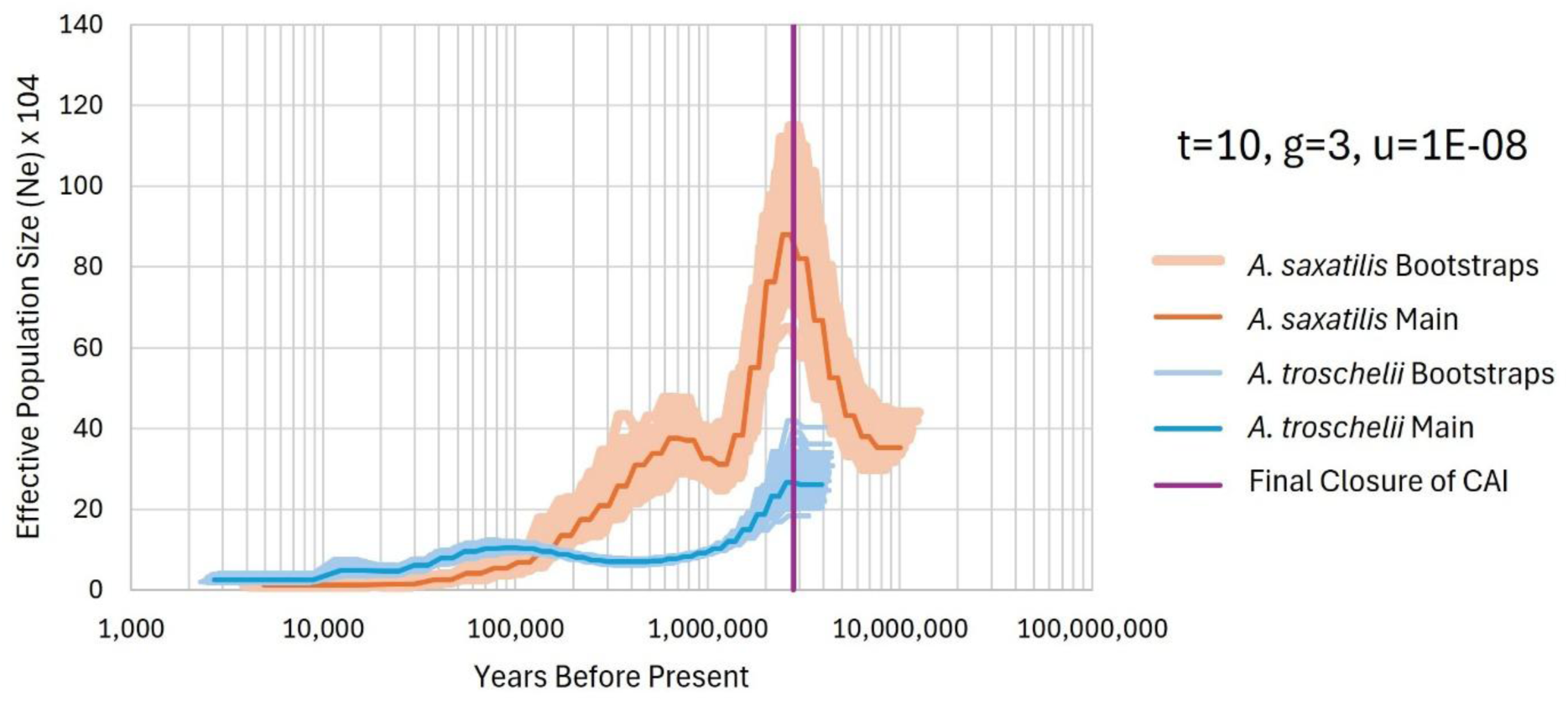
PSMC estimates of effective population size of the genomes of A. saxatilis (orange) and A. troschelii (blue). Dark lines indicate the average posterior estimate, while lighter colored lines indicate a bootstrap estimate. The purple vertical line represents the time of final closure of the CAI at 2.8 MYA.

The PSMC analysis identified a peak in the effective population size (Ne) of both *A. saxatilis* and *A. troschelii* around 2.8 MYA. This was followed by a decline in effective population size for *A. troschelii,* which remained very low until the limit of the resolution of our analysis approximately 3 thousand years ago (KYA, Figure 3). Meanwhile, *A. saxatilis* showed a decline, followed by a subsequent increase that occurred around 700 KYA, which was followed by a decline until the limit of the resolution of our analyses. Overall, the estimates showed a trend of larger population size of *A. troschelii* than *A. saxatilis* beginning at 100 KYA. Similar results were found in the shape of estimates when PSMC was run with different parameters, however the timing and values for effective population size varied significantly when changing parameters such as mutation rate (u) and generation time (g) for the species, or PSMC parameters such as -t or -p (Figures S4-S6).

The increase and subsequent decline in effective population size for both species could be associated with the oceanographic changes experienced in the TWA and the TEP following the closure of the CAI. Both species also had an overall plateau or slight decline in effective population size the past 100 KY. This is similar to patterns observed in intertidal limpets (Giles et al., 2025) but it contrasts with recent studies in fishes that have documented rapid expansions occurring after the last glacial maximum (Gatins, 2021). Furthermore, the low estimates of effective population sizes from present back to 100 KYA are unexpected, as these species are currently abundant in reefs throughout the Caribbean and Pacific (Froese & D. Pauly, 2024). It is possible that these methods lack the resolution to show more recent changes in effective population size after the last glacial maximum (Nadachowska-Brzyska et al., 2016). Further, PSMC results are likely to reflect variation in not just effective population size but also population genetic structure and selection regimes in the two sides of the CAI since these two processes are also expected to affect coalescence times (Mather et al., 2020).

### Analysis of gene family expansion

An analysis of gene family size was conducted using CAFE (Mendes et al., 2020) to identify gene families that have expanded or contracted in the two species. This analysis found 118 gene families significantly expanded or contracted in *A. saxatilis*, 104 in *A. troschelii*, and 63 in their most recent common ancestor. Among the gene families significantly expanded/contracted in *A. saxatilis* and *A. troschelii*, 41 were significant in both species.

Many of the gene families that were significantly expanded or contracted were expected due to their variability among taxa, such as genes associated with immunity. Nevertheless, a few functional groupings stood out as candidates for adaptation to the respective environments of the TWA and TEP, such as vision, reproduction, olfaction, stress response, and biosynthetic pathways. One gene family that plays a role in vision is gamma crystallin M, which is involved in eye lens development (Bloemendal et al., 2004). The expansion of gamma crystallin M in *A. saxatilis* (41 genes) and contraction in *A. troschelii* (27 genes; Figure 4) may be an indicator of their evolution in different light environments. The TWA has clearer water due to lower productivity and fewer nutrients than the TEP, and gene families influencing vision may be under greater selection in clear-water environments compared to the turbid waters of the Pacific.

**Fig. 4.**
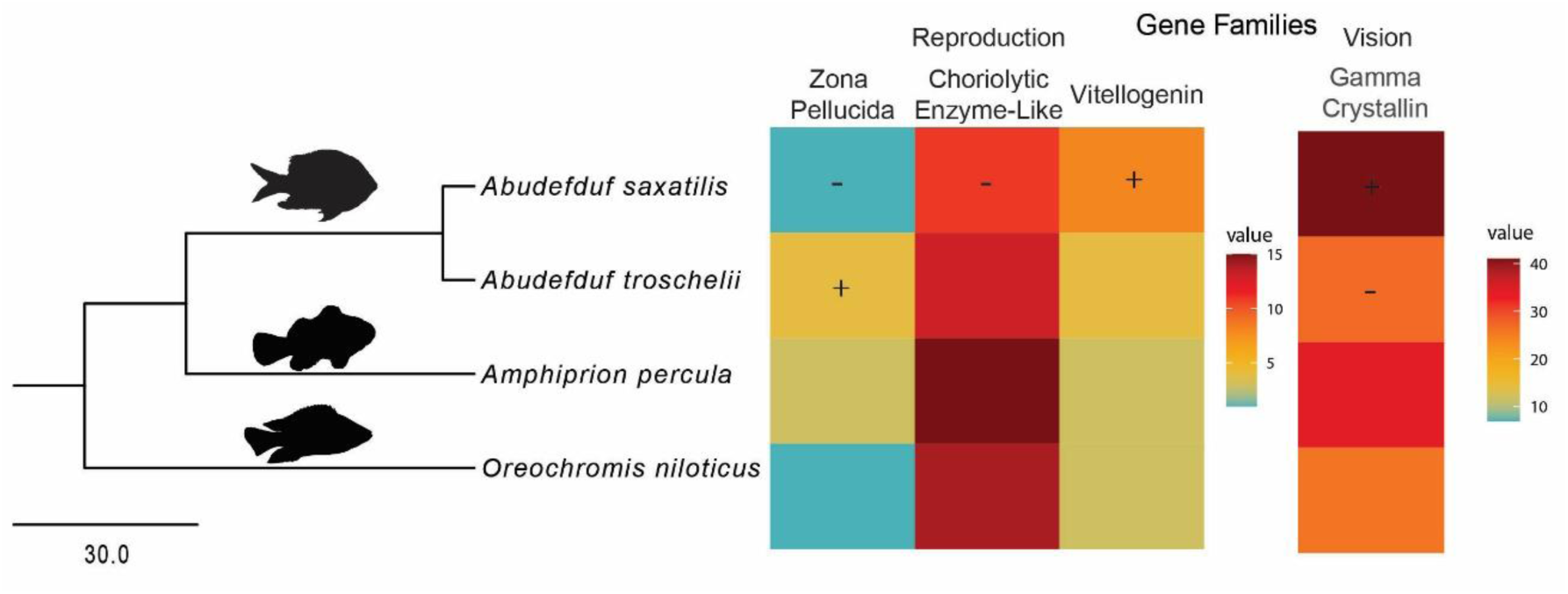
Gene family expansion of genes related to reproduction (zona pelucida, high and low choriolytic enzyme-like, and vitellogenin) and vision (gamma crystalin M family) across the phylogeny. Significant expansions and contractions are indicated with a + and –, respectively.

Three findings could be associated with reproductive differences between the transisthmian species. Vitellogenins (*vtg*) are the dominant contributors to vertebrate egg yolk and were significantly expanded in *A. saxatilis* (8 genes) compared to *A. troschelii* (4 genes). This expansion of proteins may indicate increased resources to egg development, which is consistent with previous findings that *A. saxatilis* has higher nutritional provisioning to hatchlings and a faster growth rate than *A. troschelii* (Wellington & Robertson, 2001). This is consistent with the regional-productivity hypothesis demonstrating that Caribbean species produce larger eggs than their close relatives in the Pacific due to lower larval food supplies in the Caribbean (Jackson & O’Dea, 2023; Robertson & Collin, 2015). In a similar vein, a zona pellucida protein gene family (*zpax4*), which plays a role in forming the egg casing, was significantly expanded in *A. troschelii* (4 genes) and contracted in *A. saxatilis* (1 gene). Zona pellucida genes have been shown to vary across the fish phylogeny and multiple gene duplication events have been previously identified (Sano et al., 2022). High and low choriolytic enzyme-like (*hce*), which are the two types of hatching enzymes which cooperatively digest the egg envelope (Kawaguchi et al., 2006, 2015), were also significantly contracted in *A. saxatilis* (11 genes) compared to *A. troschelii* (13 genes). Thus, two gene families that relate to egg casing formation and digestion, respectively, are contracted in *A. saxatilis* compared to *A. troschelii*.

While these gene family expansions and contractions show an interesting tie to the environment in which the species evolved, there are important caveats. First, while the gene families discussed were significantly expanded or contracted between this transisthmian species pair, this is not an exhaustive search of these gene types within the genomes. Secondly, this analysis relies on methods of gene family clustering that can lead to false identification of orthologs, and this is susceptible to different results based on the input parameters and clustering method employed. Finally, while we were able to identify significant expansions and contractions of gene families and speculate on how that may be significant given the species’ environment, additional analyses are needed to link these expansions and contractions to functional changes between these species. With this in mind, we recommend future studies that can directly link the hypothesized expanded gene families with the environmental differences in the TWA and TEP.

## Conclusion

Despite the rich history of applying molecular tools to the study of species separated by the CAI, this manuscript represents the first effort to characterize the genomic divergence of a transisthmian species pair in the TWA and TEP. The analyses presented here showed high conservation of the genome structure between transisthmian *Abudefduf*, with three large inversions between the two species. These inversions might have played a role in adaptation and speciation in *Abudefduf*, but further analyses with broader sampling are necessary to test this hypothesis. Many gene families that were significantly expanded or contracted are seemingly a result of adaptation to specific environmental conditions and the development of reproductive isolation. These findings, particularly the expansion of vitellogenin in *A. saxatilis*, are consistent with previous findings and hypotheses of differential resource allocation to egg development among species in the TEP and TWA. These genomes serve as resources for future studies on geminate species pairs, to improve our understanding of the evolutionary processes promoted by the closure of the CAI.

## Materials and Methods

### Sampling

One individual of *A. saxatilis* was obtained from an aquarium distributor (www.saltwaterfish.com) in July 2020, through whom the fish had been collected from Florida. The fish was shipped to the laboratory and was euthanized via cervical transection and tissue samples dissected (Auburn University IACUC Protocol Number PRN 2020-3708).

One adult individual of *A. troschelii* was collected on Isla Cocos (Coiba, Panama) in November 2020 using a pole spear. Muscle tissue was immediately dissected and fast frozen on liquid nitrogen. Upon arrival to Panama City tissue sample was transferred to −80°C for storage at the Smithsonian Tropical Research Institute Collection (Mi Ambiente collection permit: SE/AO-4-19).

### Genome sequencing

Gill and muscle tissue of one individual of *A. saxatilis* was dissected and immediately used for extraction of genomic DNA using the Qiagen Genomic-tip 500/G Kit. The genomic DNA extracts were sent to Novogene for PacBio Sequel II DNA HiFi library preparation and sequencing using a PacBio Sequel II system (HiFi/CCS mode) with 2 flow cells.

For *A. troschelii,* high molecular weight DNA was extracted from frozen muscle tissue using a phenol:chloroform method (Sambrook & Russell, 2006). The DNA was cleaned with 3X KAPA pure Beads (Roche Sequencing) and concentration and purity of the extracted DNA was assessed using a Qubit fluorometer (Thermo Fisher Scientific). The integrity of the DNA was evaluated with a field inversion gel (Pippin Pulse, Sage Science). On average, DNA fragments were ~50 kb in length. The PacBio HiFi library was prepared with the SMRTbell Express Template Prep Kit 2.0 (PacBio 100-938-900) using the PacBio low input protocol (DNA sheared to 15 kb) for HiFi sequencing followed by AMPure bead cleanup. The genomic library was sequenced on a single 30-h movie 8M SMRT cells in CCS mode on the PacBio Sequel II system at the Brigham Young University (BYU) DNA Sequencing Center.

### Genome de novo assembly and annotation

For both species, the raw genomic reads were assembled de novo using PacBio’s assembler HifiASM v. 0.16.1 (Cheng et al., 2021). Contamination in the genome was identified using blobtools v. 3.1.0 (Challis et al., 2020; Laetsch & Blaxter, 2017) and removed prior to annotation and comparative analyses. The completeness of the assembled genome was evaluated using BUSCO v. 5 Actinoptyergii dataset (Simão et al., 2015). Heterozygosity of the raw reads was determined by counting k-mers in Jellyfish (Marçais & Kingsford, 2011) and calculating heterozygosity in GenomeScope (Vurture et al., 2017). Reads were mapped back to the assembled genome to examine completeness using Merqury (Rhie et al., 2020). Genome annotation was conducted with ESGA (Torres & Höppner, 2021). This pipeline identifies and masks repeats using RepeatModeler v. 2.0.2 (Flynn et al., 2020) and RepeatMasker v. 4.1.2 (Smit et al., 2013) and then proceeds with Augustus v. 3.4.0 (Stanke et al., 2008) for gene model identification and training. An assembled transcriptome of *A. saxatilis* was also provided to Augustus to build gene models (Swank et al., 2024). The third dataset provided to this pipeline were known protein data from three species of the Pomacentridae family: *Acanthochromis polyacanthus*, *Amphiprion ocellaris*, and *Stegastes partitus*. Finally, this pipeline uses EVidenceModeler v. 1.1.1 (Haas et al., 2008) and PASA v. 2.5.1 (Haas et al., 2003) to build consensus gene models. The consensus gene models were functionally annotated using eggNOG-mapper v2.1.9 (Cantalapiedra et al., 2021), based on eggNOG orthology data (Huerta-Cepas et al., 2019), with the flag --tax_scope 7898 to restrict the taxonomic scope used for annotation to Actinopterygii.

### Comparative genomic analyses

A pairwise comparison of synteny was conducted between the genomes of *A. saxatilis* and *A. troschelii* to visualize structural variation in the genomes of the geminate sister species. Synteny analyses were conducted using D-GENIES online portal (Cabanettes & Klopp, 2018) using the aligner Minimap2 v2.24 (H. Li, 2018) in D-GENIES.

Following the synteny analysis, the large inversions identified between *A. saxatilis* and *A. troschelii* (see results) was examined for the genes within and immediately surrounding the inversion. To see whether there was a disproportionate number of genes with similar GOs within the inversion, we extracted GO terms from the annotation and used these to run a rank-based gene ontology analysis using the program GO-MWU (https://github.com/z0on/GO_MWU) using a Fisher’s test (Wright et al., 2015). We tested whether specific GO terms were overrepresented in the inversion compared to the rest of the genome using all three GO categories (Molecular Function, Biological Process, and Cellular Component).

Analyses of demographic history were conducted using PSMC, which scans a genome sequentially for recombination breakpoints and, with the number of coalescence events along with estimates of mutation rate and generation time, estimates the effective population size throughout time (Nadachowska-Brzyska et al., 2016). Multiple PSMC analyses were conducted, varying the values for input parameters to examine the robustness of Ne estimates to input parameters. Specifically, they were conducted using a rate of mutation (-u) spanning from 1 × 10^−8^ to 5.97 × 10^−9^, -t values of 5 vs 10, and -p flags of “4+5*3+4”, “4+20*3+4”, and “4+25*2+4+6”, and generation times (-g) between 2-7 years. The -p and -t flags can be used together to estimate overfitting, and in all tests there was no overfitting found, so we used the more complex model with more free parameters as it provided a more detailed overview of changes in Ne over time. See the github page readme on psmc for more information on examining overfitting in the psmc output. Mutation rates used were based on estimates from previously published mutation rates for a variety of fish species. The mutation rate of Atlantic herring has been estimated to be 2.0 × 10 −9 per site per generation (Feng et al., 2017), whereas in sticklebacks the mutation rate has been estimated to be 3.7 × 10 −8 (Liu et al., 2016). A more recent study estimated the mutation rate averaged across all fishes in their study to be 5.7 × 10^−09^ using pedigree analyses (Bergeron et al., 2023), however this pushed estimates of divergence back significantly later than well-supported phylogenetic estimates. In this same study, the published estimates for *Amphiprion ocellaris* ranged from .5 × 10^−08^ to 1.2 × 10^−08^, and given the close relationship to Abudefduf, we used a mutation rate within this range of 1 × 10^−08^ as the mutation rate used in our study. Generation times ranging from 2 to 7 years were used in analyses and presented in the supplement, however we present results for a generation time of 3 years. This estimate was based on the age *A. saxatilis* reaches sexual maturity, which is ~2.5 years (Villegas-Hernández et al., 2022). In the PSMC context generation time refers to the mean age of reproducing individuals, which may be different from age at sexual maturity, and would be closer to 3 years.

A gene family expansion analysis was conducted using CAFE v5.0 (Mendes et al., 2020). A total of 17 taxa were included in the analysis to represent various clades across the fish tree of life (Table 3). Protein data for each species included was downloaded from Ensembl biomart. Proteins for each species were all compared to one another using BLAST to obtain similarity scores, and they were clustered by similarity using mcl (Van Dongen, 2008).

**Table 3.** Species included in the CAFÉ analysis, to evaluate the potential expansions or contractions of gene families among transisthmian *Abudefduf*.

| Scientific Name | Ensembl ID | Common Name | Family | Number<br>of<br>Proteins<br>Raw | # longest<br>proteins |
| --- | --- | --- | --- | --- | --- |
| *Astyanax*<br>*mexicanus* | ENSAMXG | Mexican Tetra | Characidae | 41410 | 27984 |
| *Amphiprion*<br>*percula* | ENSAPEG | Orange Clownfish | Pomacentridae | 35899 | 25045 |
| *Cottoperca*<br>*gobio* | ENSCGOG | Channel Bull Blenny | Bovichtidae | 60811 | 22935 |
| *Clupea*<br>*harengus* | ENSCHAG | Atlantic Herring | Clupeidae | 67663 | 24775 |
| *Dicentrarchus*<br>*labrax* | ENSDLAG | European Bass | Moronidae | 69565 | 23993 |
| *Danio rerio* | ENSDARG | Zebrafish | Cyprinidae | 65905 | 33856 |
| *Gasterosteus*<br>*aculeatus* | ENSGACG | Threespine<br>Stickleback | Gasterosteidae | 29245 | 21065 |
| *Gadus morhua* | ENSGMOG | Atlantic Cod | Gadidae | 68853 | 24085 |
| *Ictalurus*<br>*punctatus* | ENSIPUG | Channel Catfish | Ictaluridae | 39017 | 25196 |
| *Labrus bergylta* | ENSLBEG | Ballen Wrasse | Labridae | 40572 | 28105 |

| Scientific Name | Ensembl ID | Common Name | Family | Number of Proteins Raw | # longest proteins |
| --- | --- | --- | --- | --- | --- |
| *Larimichthys crocea* | ENSLCRG | Yellow Croaker | Sciaenidae | 66805 | 23506 |
| *Oreochromis niloticus* | ENSONIG | Nile Tilapia | Cichlidae | 82099 | 29453 |
| *Oryzias sinensis* | ENSOSIG | Chinese Rice Fish | Adrianichthyidae | 54551 | 22209 |
| *Poecilia formosa* | ENSPFOG | Amazon Molly | Poeciliidae | 31637 | 23950 |
| *Salmo salar* | ENSSSAG | Atlantic Salmon | Salmonidae | 184209 | 51777 |
| *Abudefduf saxatilis* | NA | Sergeant Major<br>Damsel fish | Pomacentridae | NA | 25888 |
| *Abudefduf troschelii* | NA | Panamanian Sergeant<br>Major | Pomacentridae | NA | 25261 |

A time-calibrated phylogenetic tree was created by downloading sequences used in the fish tree of life (Chang et al., 2019; Rabosky et al., 2018) and subsampling it to only include the taxa used in the CAFE analysis. This alignment was then edited manually to remove loci for which data was missing for >58% of the taxa (removed genes with 7 or fewer samples). The tree was created using BEAST v2.5 (Bouckaert et al., 2019) with bmodeltest (Bouckaert & Drummond, 2017) to infer the model of evolution for each locus and was run for 50,000,000 generations sampling the posterior every 5,000 generation. Two fossil calibrations used in the Fish Tree of Life that applied to the nodes in the subset tree were used. These are on the most recent common ancestor (MRCA) of all Pomacentridae with a gamma distribution offset 44 and an alpha and beta parameter of 5 and 3 respectively (including *Abudefduf saxatilis, Abudefduf troschelii,* and *Amphiprion percula*), and the second on the MRCA of Otomorpha with a gamma distribution offset 145, with an alpha parameter of 5 and a beta parameter of 3 (including *Astyanax mexicanus, Clupea harengus, Danio rerio, and Ictalurus punctatus*).

## Data Availability

Raw reads are available through NCBI SRA (BioProject: PRJNA1244908, Temporary Submission ID: SUB15182265, *A. saxatilis* BioSample: SAMN47735137 and read ID: SRR32938816, *A. troschelii* BioSample: SAMN47735138 and read ID: SRR32938815). Scripts used for analyses are available on github: https://github.com/C-Tracy/GenomeAlignmentAnnotation.

## Acknowledgements

We would like to thank Adam Hallaj for his assistance with DNA extractions, Katie Eaton for her helpful guidance on analyses, and the Bernal Lab and the Phyletica Lab and specifically Matthew Buehler for discussions and feedback on the project. Computational work was possible through resources provided by the Easley Computing Cluster at Auburn University.

