## Supplemental Tables and Figures for "Comparative Genomics of Transisthmian Damselfishes (*Abudefduf saxatilis* and *A. troschelii*)"

### Supplemental Figures

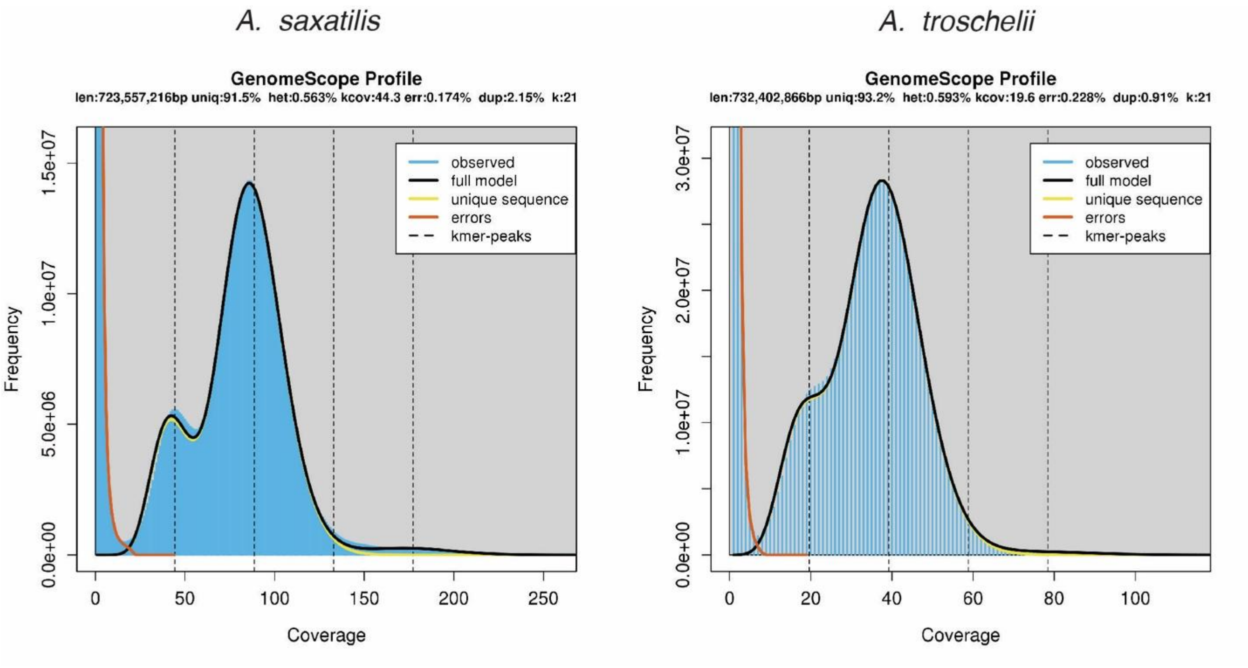


**Figure S1.** Frequency and coverage of the genome of *A. saxatilis* (left) and *A. troschelii* (right) from GenomeScope of the k-mer count and distribution determined using Jellyfish. The differences in coverage between the two species are from the sequencing efforts and are not associated with biological differences (see methods for more details).

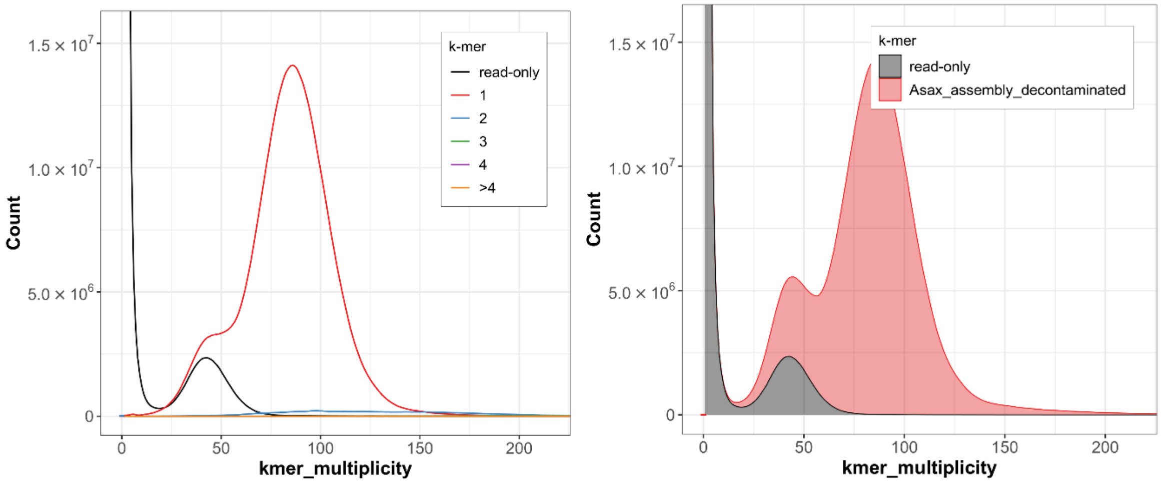


**Figure S2.** Merqury results for *A. saxatilis* showing how often k-mers from the reads are found in the assembly (left), and the distribution of k-mers from reads and assembly with their multiplicity in the reads (right).

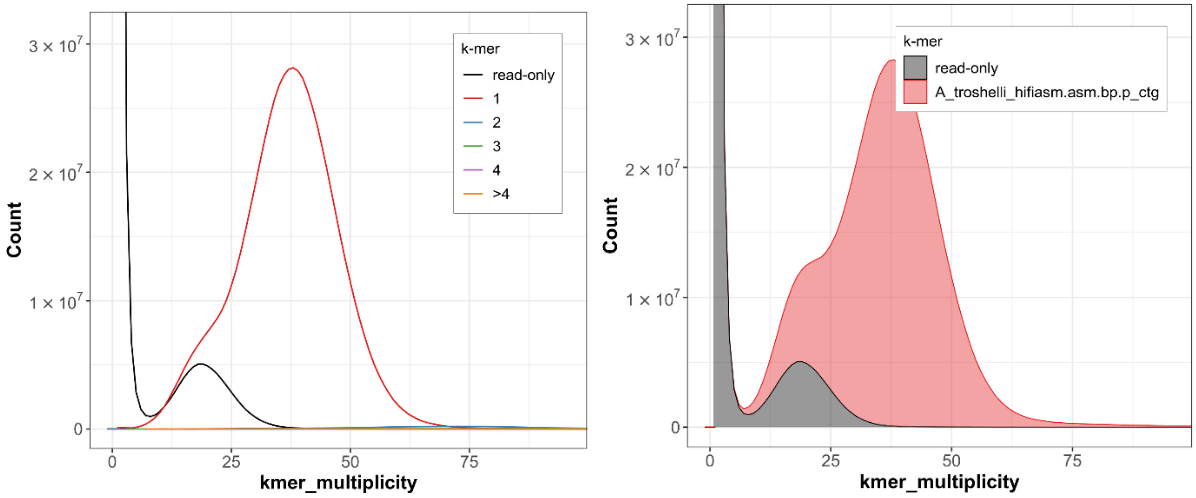


**Figure S3.** Merqury results for *A. troschelii* showing how often k-mers from the reads are found in the assembly (left), and the distribution of k-mers from reads and assembly with their multiplicity in the reads (right).

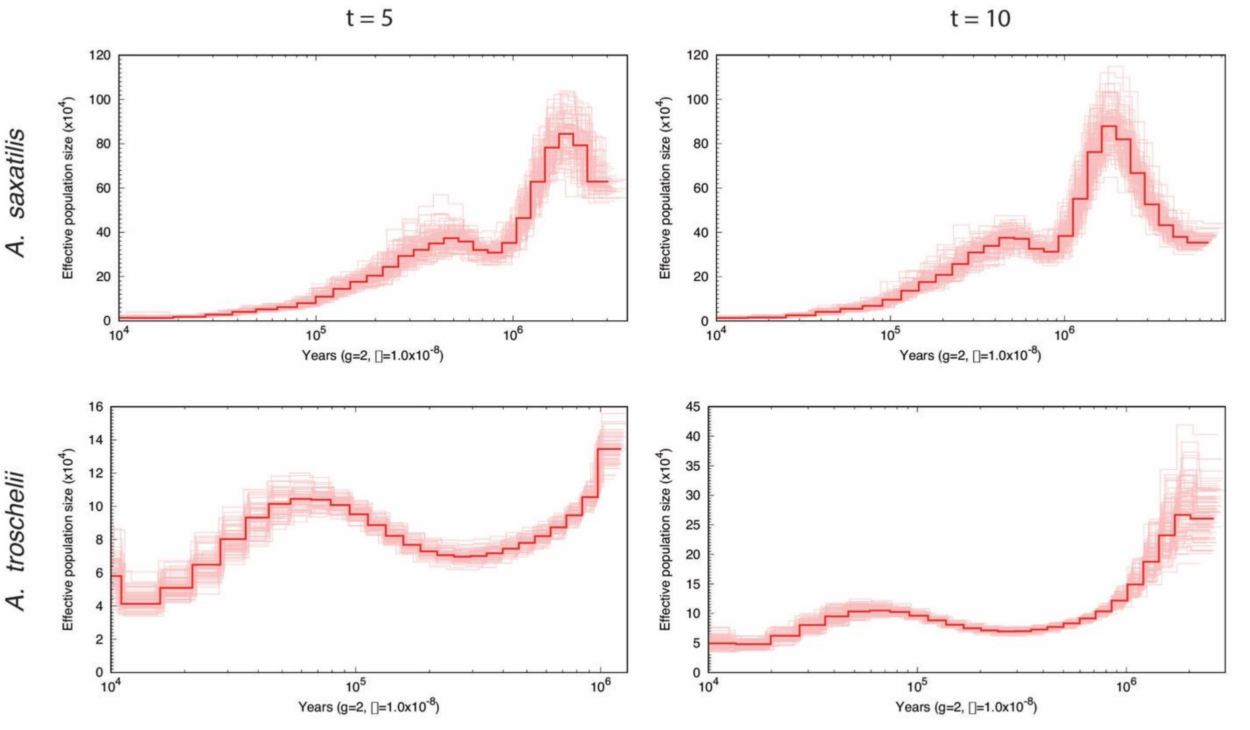


**Figure S4.** Demographic history results from PSMC run for *A. saxatilis* (top) and *A. troschelii* (bottom) run with t = 5 (left) and t = 10 (right) run with 100 bootstrap replicates in each analysis. Dark red is the result from a consensus estimate, and all the light pink lines are the results from each bootstrap run. All graphs were created with an estimated generation time of 2 years. Analyses were run several times with different parameters, however the pattern of the curve did not change for either species.

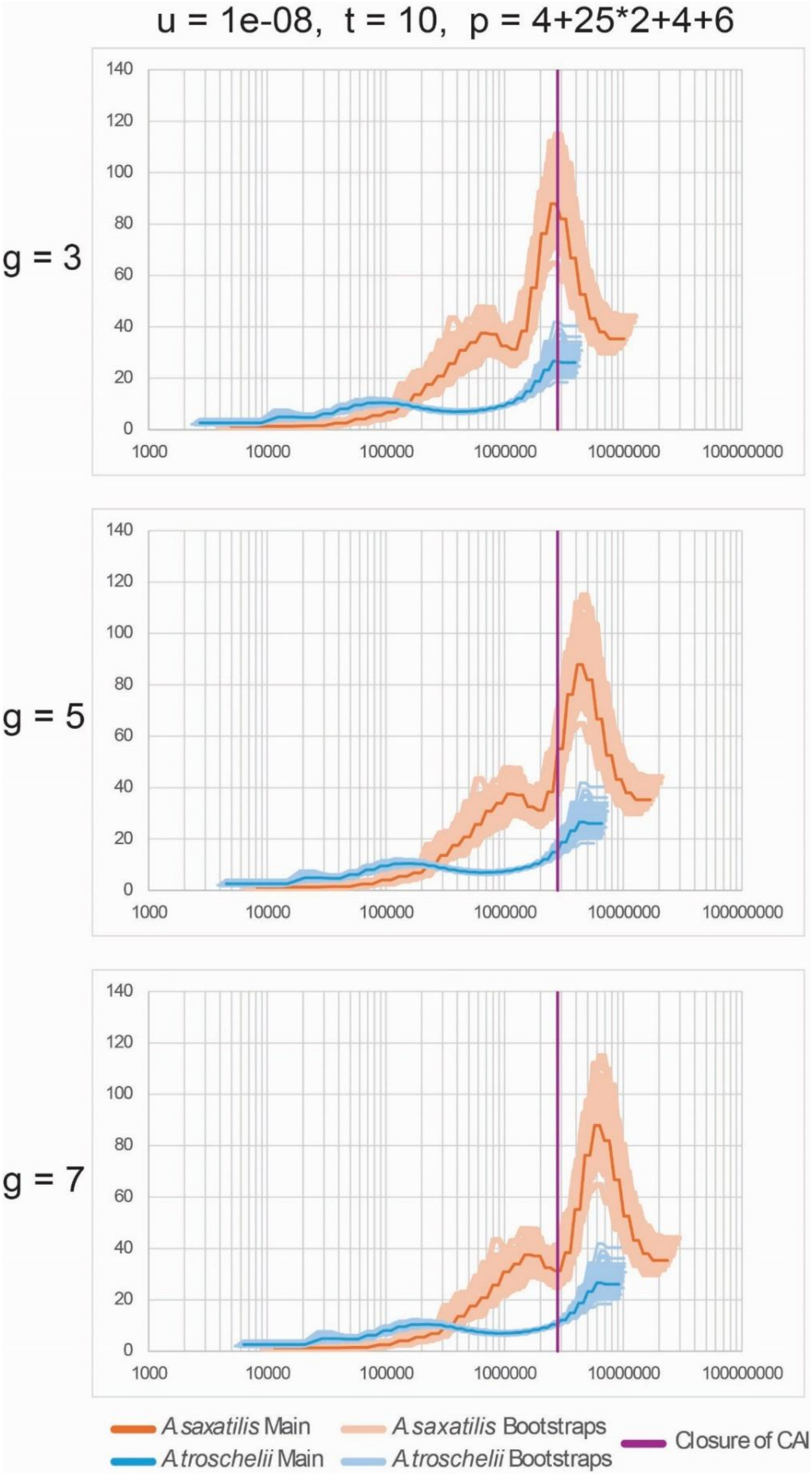


**Figure S5.** Estimates of effective population size with PSMC, changing the generation time and keeping all other parameters constant.

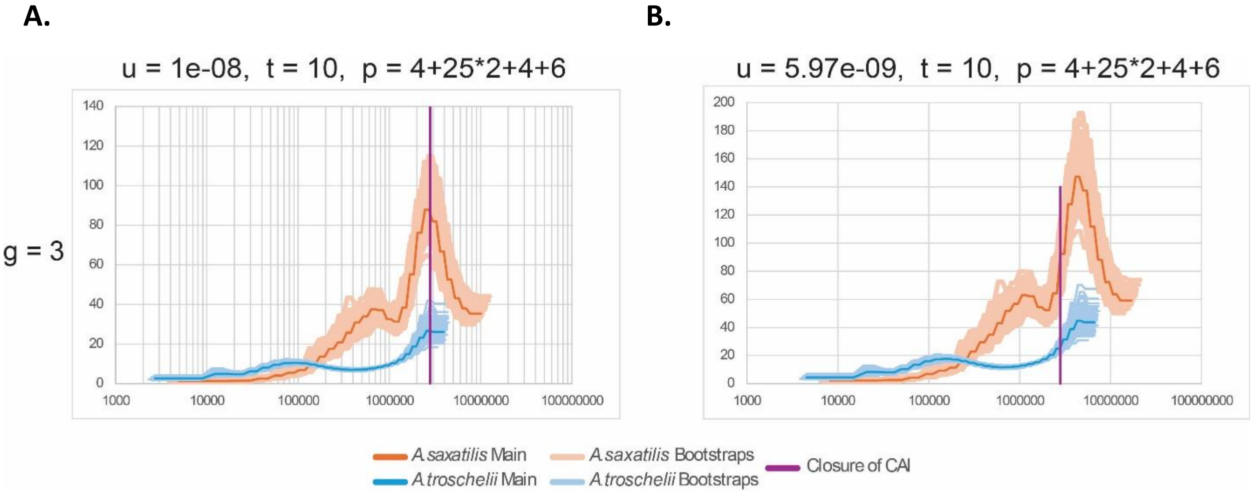


**Figure S6.** Estimates of effective population size with PSMC, changing the mutation rate on and keeping all other parameters constant. Mutation rate on panel A is within the range published for *Amphiprion ocellaris* in Bergeron et al., 2023, while the mutation rate in panel B is the average for all fishes from Bergeron et al., 2023.

**Table S1.** Mapping of *A. saxatilis* contigs to Nemo genome assembly chromosomes. Contigs highlighted in red are mapped to multiple Nemo contigs, and contigs in green are mapped to only one Nemo contig.

| Nemo Contig ID | Nemo Chromosome Number | Percent Mismatch | Number mapped reads | Features | A.saxatilis mapped contigs |
|---|---|---|---|---|---|
| CM009708 | 1 | 2.9 | 4 | 0 | ptg000023l<br>ptg000040l<br>ptg000120l<br>ptg000138l |
| CM009709 | 2 | 3 | 3 | 0 | ptg000007l<br>ptg000034l<br>ptg000046l |
| CM009710 | 3 | 4.2 | 9 | 0 | ptg000027l<br>ptg000029l<br>ptg000062l<br>ptg000026l<br>ptg000067l<br>ptg000025l<br>ptg000034l<br>ptg000053l<br>ptg000092l |
| CM009711 | 4 | 3.2 | 4 | 0 | ptg000008l<br>ptg000015l<br>ptg000033l<br>ptg000053l |
| CM009712 | 5 | 3.1 | 2 | 0 | ptg000014l<br>ptg000045l |
| CM009713 | 6 | 4.5 | 8 | 0 | ptg000072l<br>ptg000040l<br>ptg000067l<br>ptg000026l<br>ptg000034l<br>ptg000053l<br>ptg000039l<br>ptg000077l |
| | 7 | 2.9 | 4 | 0 | ptg000002l |

| | | | | | |
|---|---|---|---|---|---|
| CM009714 | | | | | ptg0000211<br>ptg0000301<br>ptg0000771 |
| CM009715 | 8 | 3.1 | 2 | 0 | ptg0000361<br>ptg0000391 |
| CM009716 | 9 | 4.7 | 15 | 0 | ptg0000051<br>ptg0000191<br>ptg0000291<br>ptg0000161<br>ptg0000391<br>ptg0000531<br>ptg0000411<br>ptg0000321<br>ptg0000041<br>ptg0000581<br>ptg0000761<br>ptg0000621<br>ptg0000921<br>ptg0000381<br>ptg0000811 |
| CM009717 | 10 | 3.5 | 2 | 0 | ptg0000111<br>ptg0000281 |
| CM009718 | 11 | 3 | 3 | 0 | ptg0000061<br>ptg0000311<br>ptg0000381 |
| CM009719 | 12 | 2.8 | 2 | 0 | ptg0000041<br>ptg0000581 |
| CM009720 | 13 | 6.5 | 14 | 2 | ptg0000191<br>ptg0000621<br>ptg0000251<br>ptg0000291<br>ptg0000921<br>ptg0000261<br>ptg0000671<br>ptg0000541<br>ptg0000321<br>ptg0000771<br>ptg0000341<br>ptg0000181 |

| | | | | | |
|---|---|---|---|---|---|
| | | | | | ptg000072l<br>ptg000003l |
| CM00972<br>1 | 14 | 3.3 | 3 | 0 | ptg000001l<br>ptg000013l<br>ptg000035l |
| CM00972<br>2 | 15 | 4.1 | 16 | 0 | ptg000018l<br>ptg000043l<br>ptg000024l<br>ptg000038l<br>ptg000092l<br>ptg000062l<br>ptg000016l<br>ptg000076l<br>ptg000053l<br>ptg000026l<br>ptg000058l<br>ptg000004l<br>ptg000032l<br>ptg000041l<br>ptg000039l<br>ptg000005l |
| CM00972<br>3 | 16 | 67.4 | 31 | 281 | ptg000009l<br>ptg000044l<br>ptg000038l<br>ptg000129l<br>ptg000173l<br>ptg000226l<br>ptg000271l<br>ptg000151l<br>ptg000166l<br>ptg000177l<br>ptg000192l<br>ptg000197l<br>ptg000227l<br>ptg000235l<br>ptg000259c<br>ptg000154c<br>ptg000248l<br>ptg000141l |

| | | | | | |
|---|---|---|---|---|---|
| | | | | | ptg000268l<br>ptg000027l<br>ptg000182l<br>ptg000053l<br>ptg000039l<br>ptg000025l<br>ptg000092l<br>ptg000029l<br>ptg000062l<br>ptg000019l<br>ptg000026l<br>ptg000067l<br>ptg000088l |
| CM009724 | 17 | 3.6 | 1 | 0 | ptg000010l |
| CM009725 | 18 | 3 | 3 | 0 | ptg000016l<br>ptg000020l<br>ptg000034l |
| CM009726 | 19 | 2.9 | 2 | 0 | ptg000017l<br>ptg000107l |
| CM009727 | 20 | 3 | 3 | 0 | ptg000012l<br>ptg000035l<br>ptg000080l |
| CM009728 | 21 | 2.9 | 2 | 0 | ptg000008l<br>ptg000050l |
| CM009729 | 22 | 3 | 3 | 0 | ptg000003l<br>ptg000041l<br>ptg000065l |
| CM009730 | 23 | 4.1 | 13 | 0 | ptg000019l<br>ptg000016l<br>ptg000005l<br>ptg000039l<br>ptg000053l<br>ptg000041l<br>ptg000032l<br>ptg000004l<br>ptg000058l<br>ptg000076l<br>ptg000062l<br>ptg000092l |

| | | | | | |
|---|---|---|---|---|---|
| | | | | | ptg000038l |
| CM00973 1 | 24 | 28 | 23 | 1 | ptg000062l<br>ptg000026l<br>ptg000025l<br>ptg000077l<br>ptg000003l<br>ptg000034l<br>ptg000019l<br>ptg000072l<br>ptg000018l<br>ptg000040l<br>ptg000067l<br>ptg000016l<br>ptg000005l<br>ptg000039l<br>ptg000053l<br>ptg000041l<br>ptg000032l<br>ptg000092l<br>ptg000004l<br>ptg000058l<br>ptg000076l<br>ptg000038l<br>ptg000024l |
